# ProAffinity++

**DOI:** 10.1101/2025.10.31.685718

**Authors:** Keyu Xu, Zeyuan Di, Jianquan Zhao, Haicang Zhang, Dongbo Bu

## Abstract

Proteins are essential biological macromolecules that play a crucial role in living organisms. Protein-protein interactions, which govern various biological processes such as signal transduction, cell metabolism, and cell growth, are key aspect of protein function. The strength of these interactions, characterized by protein-protein binding affinity(**Δ**G), is a critical thermodynamic property of protein complexes. Accurate prediction of protein-protein binding affinity is meaningful for understanding the mechanisms of biological systems and improving the speed of binding affinity determination. However, current methods are limited in their accuracy and are primarily designed for binary complexes. To address these limitations, we propose ProAffinity++, an end-to-end algorithm that predicts binding affinities using a combined representation of sequence and structure. The core idea of ProAffinity++ is to model the binding regions of protein complexes and utilize graph neural networks to capture the local microenvironment of residues. Our experimental evaluations demonstrate that ProAffinity++ outperforms state-of-the-art methods in predicting protein-protein binding affinities. Moreover, it exhibits remarkable performance in challenging scenarios, such as antigen-antibody interactions and missense mutation problems, where existing methods have limitations.

## 1 Introduction

In protein science, binding affinity plays a pivotal role in various domains[1, 2]. For instance, in immunotherapy for cancer treatment, T cells recognize and attack cancer cells by binding the complementarity determining regions (CDRs) of T cell receptors (TCRs) to peptides presented by major histocompatibility complexes (MHCs) on cancer cells[3]. In this way, accurately determining binding affinity is essential for designing TCRs that effectively recognize and target cancer cells, thereby enhancing the efficacy of cancer treatment.

To measure binding affinities, biologists have developed various experimental methods, including surface plasmon resonance (SPR)[4] and isothermal titration calorimetry (ITC)[5]. However, these methods determine protein-protein binding affinity slowly, making them unsuitable for high-throughput calculations, limiting the amount of binding affinities data.

To address the limitations of the experimental methods, various binding affinity predictors have been developed, including molecular dynamics simulations, empirical energy functions, and machine learning methods. Molecular dynamics simulations predict binding affinities accurately but cost significant computational time[6–9]. Compared to molecular dynamics simulations, empirical energy functions predict binding affinities efficiently, however, these methods decompose binding affinity into multiple energy terms manually [10–12] that may introduce biases. Machine learning methods have shown promising results in predicting binding affinity. For example, Prodigy[13] predicts binding affinity using a linear regression predictor that employs the physical and chemical properties of residues as input. PIPR[14] utilizes a convolutional neural network (CNN) to capture local features and a recurrent neural network (RNN) to retain contextual information for affinity prediction. PPI-Affinity [15] employs a support vector machine (SVM) to predict the binding affinity from the structure features that are captured by ProtDCal[16].

While several computational binding affinity predictors based on different principles have been proposed, predicting binding affinities remains challenging due to the following factors: (i) inadequate binding affinity data limited models to learn robust protein representations. (ii) the binding signal is easily diluted by other unbinding information because only the residues located at the binding interface interact with another protein; (iii) protein binding affinity is influenced by comprehensive consideration of factors.

In this paper, we introduce the deep learning algorithm ProAffinity++ to predict protein-protein binding affinity. To sufficiently capture the features of the protein complex, ProAffinity++ incorporates sequence features extracted by a pre-trained language model and structure features extracted by an inverse folding model. ProAffinity++ incorporates pre-trained language models for protein sequences and pre-trained models for protein structures to extract sequence and structure features. To obtain robust protein representations, ProAffinity++ models the binding interface of protein complexes and captures the local microenvironment of residues using a graph neural network. To make full use of protein-protein binding data, ProAffinity++ was also trained on multiple complexes.

We summarized our contributions in three aspects. (i) We developed an end-to-end binding affinity prediction algorithm that alleviates the data preprocessing burden for users. ProAffinity++ takes protein structures as input and provides options for extracting protein sequences and utilizing pre-trained models for extracting sequence and structure features. (ii) We modeled the binding regions of protein complexes and extracted the local microenvironment of residues using a graph neural network. (iii) ProAffinity++ applies to antigen-antibody, missense mutations, and multi-component protein complexes.

We evaluated our algorithm using four datasets and found that ProAffinity++ outperforms various state-of-the-art methods on the PDBBind[17] datasets and Affinity Benchmark datasets. ProAffinity++ shows strong performance on the antigen-antibody dataset AT-Bind and the missense mutation dataset SKEMPI[18] demonstrates its generalization capability.

## 2 Related works

ProAffinity++ is built upon two fundamental ideas. Firstly, ProAffinity++ leverages a comprehensive approach by extracting both sequence and structure information from proteins, thereby capturing a rich representation from protein complexes. Secondly, ProAffinity++ innovatively focuses on the binding region, rather than the entire protein complex, to predict protein-protein binding affinity.

Two methods are widely used to extract the protein sequence information. The first approach uses the Position-Specific Scoring Matrix sequence(PSSM)[19] obtained through searching multiple sequence alignments to represent the conservation of residues at different positions in the protein. Each row of the PSSM corresponds to an amino acid type, and the columns represent positions in the protein sequence, so each element in the matrix represents the amino acid score at the corresponding position. However, the PSSM heavily relies on the quality of the sequence alignment algorithm, as low-quality alignments result in inaccurate matrices. Additionally, PSSM only considers interactions between adjacent residues and ignores capturing long-range interactions. The second method directly decodes the sequence information using a protein language model like ESM-1v [20]. The self-attention mechanism enables ESM-1v to capture long-range interactions within protein sequences. ESM-1v is trained on the Uniref90[21] sequence dataset and incorporates the evolutionary information of the protein sequence by simulating sequence evolution through natural selection. ESM-1v decodes not only evolutionary information but also global information from the protein sequence.

For protein structure information, the protein inverse folding method is suitable to capture and represent the protein backbone information, because the protein inverse folding aims to recover protein sequences from protein backbone atom coordinates. ESM-IF1[22] decodes the protein backbone information based on the transformer module. To learn the decoding capability, ESM-IF1 is trained using the 12M protein structures predicted by AlphaFold2[23] and experimentally determined protein structures.

In this study, we model the binding regions of protein complexes as bipartite graphs, because the interaction patterns between different chains in protein complexes and the interactions between residues within them are bipartite graphs. The overall complex graph consists of multiple bipartite graphs, since multi-component complexes have multiple binding regions, each binding region is represented by a bipartite graph. In a bipartite graph, residues are represented as nodes, interactions between residues are represented as edges, and the chains identifiers are used for the bipartite graphs. The initial features of the bipartite graph nodes include chain identifiers, residue types, physicochemical properties, evolutionary information, side-chain information, and secondary structure information. The initial features of the edges include the distance between residue pairs and the relative position of residue pairs.

## 3 Results

### 3.1 ProAffinity++ shows better performance than other state-of-the-art methods

We compared the performance of ProAffinity++ on PDBBind Benchmark and Affinity Benchmark to other methods. As shown in the table1, the performance of the ProAffinity++ algorithm exceeded all the comparison methods, including the FoldX, Prodigy, IsLand, PPI-Affinity, PIPR, and the SeqAffinity. While ProAffinity++ trained only on the binary complex dataset S653 showed comparable performance with these methods, ProAffinity++ trained on the binary complex S653 and the multiprotein complex dataset S1902 demonstrated better performance than these methods.

**Table 1:** Performance of ProAffinity++ and state-of-the-art methods on the PDB-Bind benchmark and Affinity benchmark.

| Method | PDBBind |  |  | Affinity Benchmark |  |  |
| --- | --- | --- | --- | --- | --- | --- |
|  | Pearson | Spearman | MAE (kcal/mol) | Pearson | Spearman | MAE (kcal/mol) |
| FoldX | 0.304 | 0.521 | 6.137 | 0.472 | 0.493 | 5.387 |
| Prodigy | 0.306 | 0.473 | 2.513 | 0.735 | 0.7453 | 1.433 |
| IsLand | 0.270 | 0.264 | 2.201 | 0.378 | 0.446 | 2.101 |
| PPI-Affinity | 0.495 | 0.462 | 1.800 | 0.616 | 0.618 | 1.819 |
| PIPR | 0.516 | 0.418 | 1.745 | 0.641 | 0.634 | 1.842 |
| SeqAffinity | 0.530 | 0.533 | 1.723 | 0.716 | 0.697 | 1.452 |
| ProAffinity++ | 0.549 | 0.518 | 1.734 | 0.712 | 0.662 | 1.605 |
| **ProAffinity++(S653+S1902)** | **0.594** | **0.567** | **1.650** | **0.778** | **0.812** | **1.412** |

On PDBBind Benchmark, the Pearson correlation coefficient of the ProAffinity++ method reached 0.676, which was 0.064 higher than the best performance of state-of-the-art methods the Spearman correlation coefficient was 0.034 higher than the best performance of state-of-the-art methods and achieved 0.812, and the absolute mean error(MAE) was decreased by 0.073 kcal/mol to 1.455 kcal/mol. On Affinity Benchmark, the ProAffinity++ also manifested consistent results the Pearson correlation coefficient of reached 0.778, the Spearman correlation coefficient reached 0.635, and the absolute mean error(MAE) decreased to 1.412 kcal/mol.

The predicted Δ*G* of ProAffinity++ is significantly correlated with the realized Δ*G*. These results indicate that ProAffinity++ can capture the intrinsic mechanism of the binding affinity of protein complexes.

### 3.2 ProAffinity++ is useful in predicting the change of binding affinity for nonsynonymous substitution

**Table 2:**
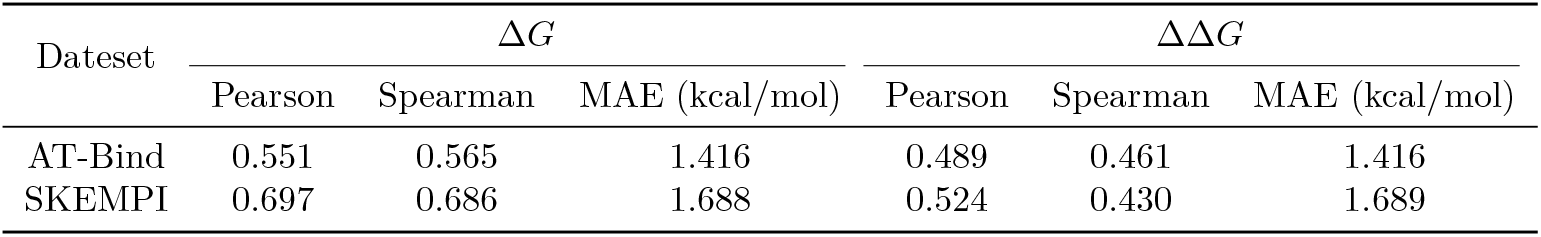
Performance of ProAffinity++ in predicting the change of binding affinity for nonsynonymous substitution.

Predicting the change of binding affinity is useful in protein design, and drug discovery. We assume that the binding affinity can be simply calculated by the mutated binding affinity subtracting the wildtype binding affinity. ProAffinity++ exhibited outstanding performance in protein-protein binding affinity prediction, so we tried to apply ProAffinity++ in calculating the protein-protein binding affinity changes for the nonsynonymous substitution according to our assumption.

The performance of ProAffinity++ in protein-protein binding affinity change prediction on the AT-Bind and the SKEMPI dataset was shown in the table2 the predicted ΔΔ*G* and the ground-truth ΔΔ*G* correlation was shown in the figure1. Although ProAffinity++ achieved almost meaningful Pearson correlation coefficient and Spearman correlation coefficient on the AT-Bind and the SKEMPI dataset, we found that the performance of ProAffinity++ in protein-protein binding affinity change prediction was lower than the performance of ProAffinity++ in protein-protein binding affinity prediction.

**Fig. 1:**
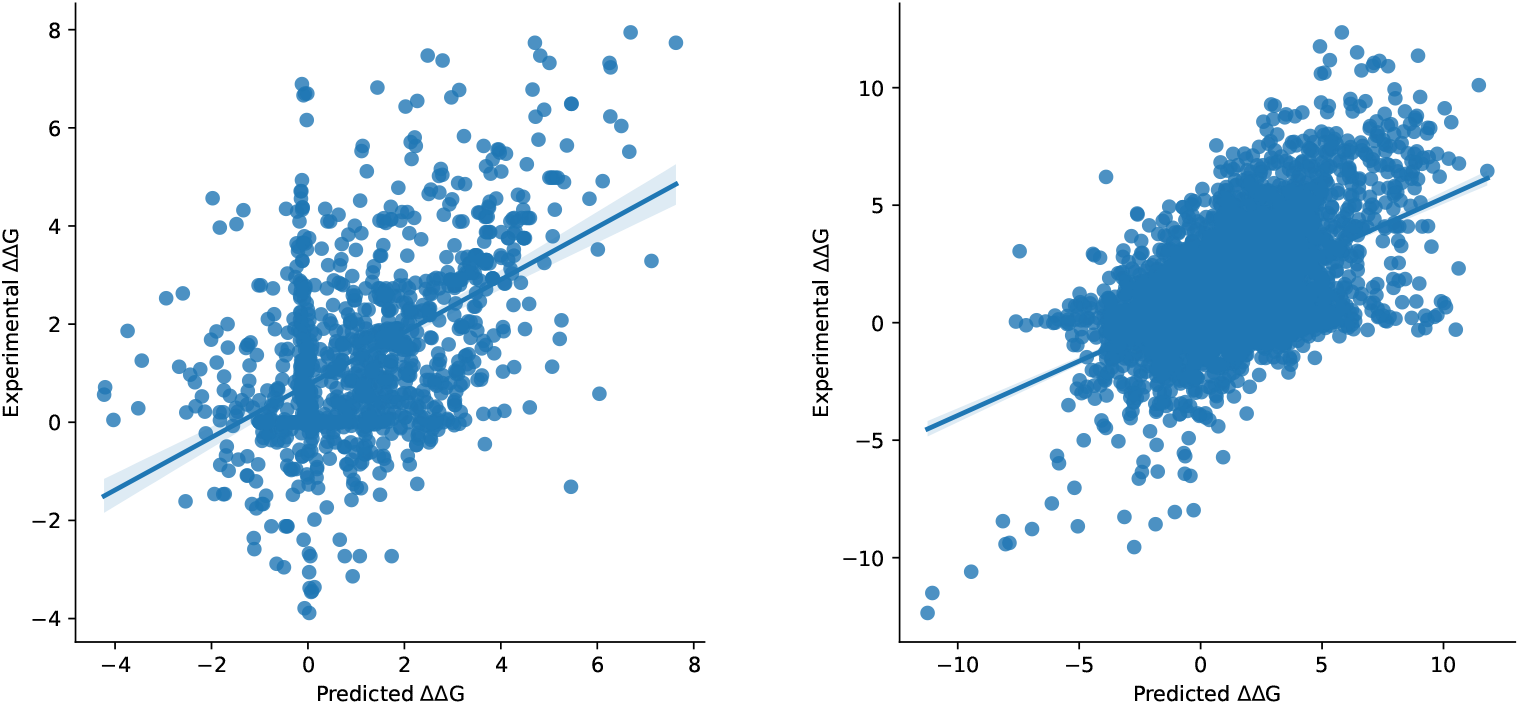
The performance of ProAffinity++ in delta delta G

**Fig. 2:**
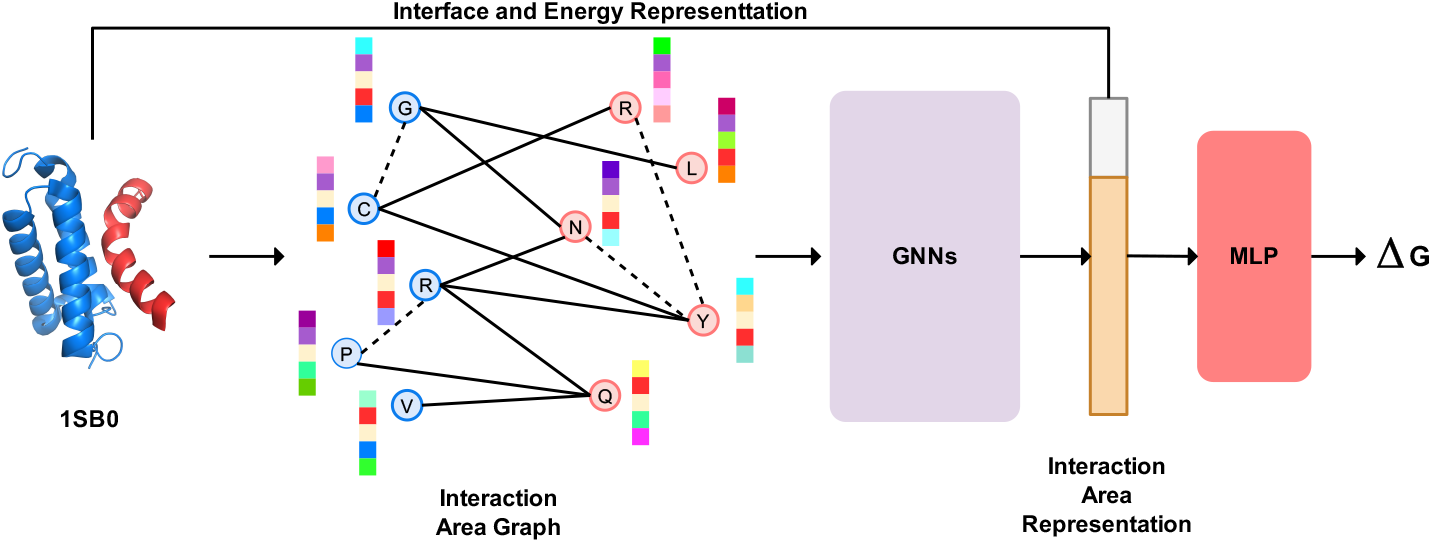
ProAffinity++ overview. ProAffinity++ first built the interaction area graph through a bipartite graph according to the protein structure. Then, ProAffinity++ used a GNN to update the node representations of the initial bipartite graph. Finally, a multilayer perceptron predicted the binding affinity based on the interaction area representation and the physical energy representation.

**Fig. 3:**
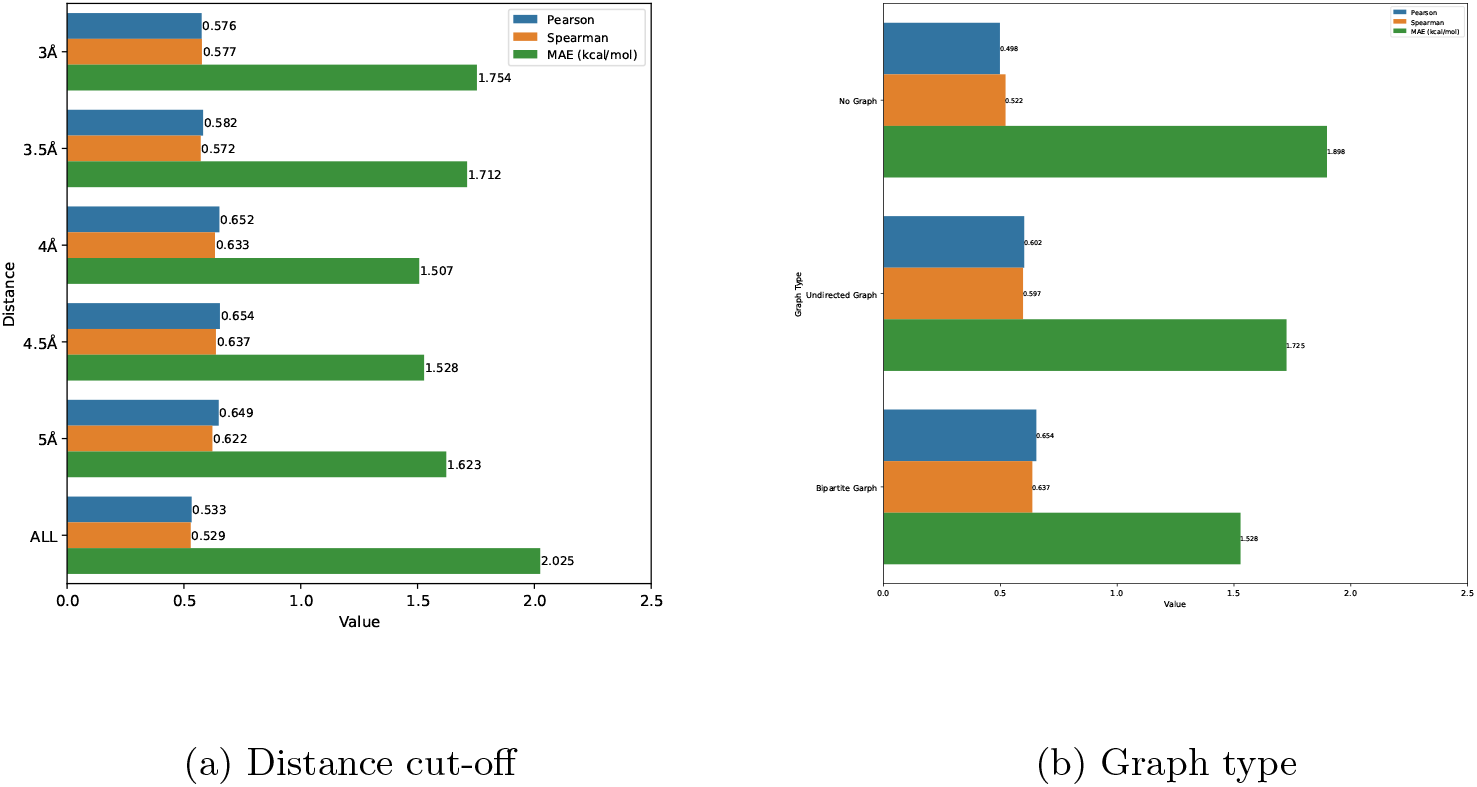
The performance of ProAffinity++ with different graph types and distance cut-offs. (a) Keeping partial residue pairs for building a bipartite graph was favorable for the performance of ProAffinity++ while keeping all residue pairs significantly reduced the performance of ProAffinity++. (b) The application of a bipartite graph lifted the Performance of ProAffinity++ compared to an undirected graph or removed the graph module.

This result demonstrated that the binding affinity change is not as simple as we assumed, in other words, the binding affinity change does not obey the base linear operations, a more precise binding affinity change predictor should consider more comprehensive factors.

### 3.3 ESM-1v representation significantly improves the performance of ProAffinity++

**Table 3:** Ablation of modules of ProAffinity++.

| Module | Pearson | Spearman | MAE (kcal/mol) |
| --- | --- | --- | --- |
| ProAffinity++ | 0.654 | 0.637 | 1.528 |
| w/o AAindex | 0.631 | 0.611 | 1.643 |
| w/o SideChain | 0.587 | 0.592 | 1.744 |
| w/o DSSP | 0.601 | 0.587 | 1.697 |
| w/o ESM-1v | 0.496 | 0.441 | 2.043 |
| w/o ESM-IF1 | 0.584 | 0.581 | 1.741 |
| w/o FoldX | 0.551 | 0.489 | 1.785 |
| w/o ASA | 0.645 | 0.614 | 1.552 |
| w/o Distance | 0.621 | 0.601 | 1.685 |

ProAffinity++ combined different level features including the sidechain information, protein sequence representation using the ESM-1v, protein structure information captured by ESM-IF, physical energy information calculated by FoldX, and other information. To analyze the importance of the different features, we conducted an ablation experiment for the PreAffinity++ through 5-fold validation on the training dataset(Sup. A.1), the ablation result was presented in the table3.

According to the principle that the protein sequence decides the protein structure and the protein structure decides the protein function, the ablation study reveals that the protein sequence representation profoundly impacts the predictive performance of ProAffinity++, suggesting that ESM-1v is capable of decoding diversity protein structure information from protein sequences.

We also found that the energy terms of the FoldX played an important role in ProAffinity++. Without the FoldX module, the performance of the ProAffinity++ was reduced greatly, the Pearson correlation coefficient was reduced by 0.103, the Spearman correlation coefficient was reduced by 0.138, and the absolute average error of the binding affinity was added by 0.257 kcal/mol. FoldX fastly and quantitatively estimates the importance of the interaction of protein structure based on its force field, in this way, we believe that the physical model of the protein structure through heuristic rules corrects the unreasonable result of the neural network.

### 3.4 Adding noise and augment data improves the performance of ProAffinity++

**Table 4:**
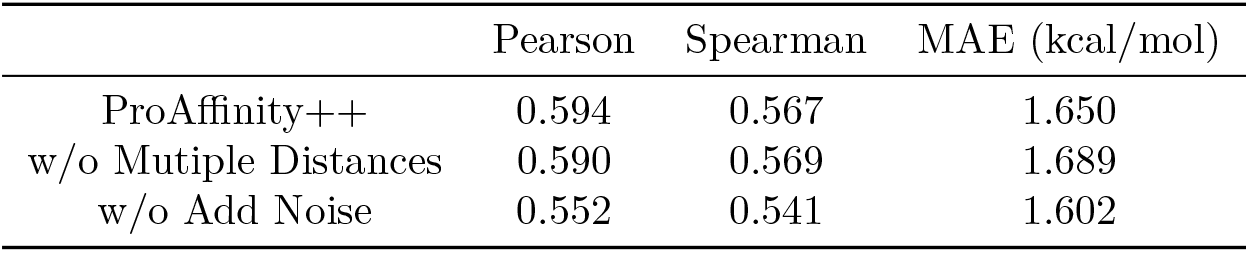
Ablation of augment method of ProAffinity++.

To enable the robustness of ProAffinity++, we injected Gaussian noise for the node features of the bipartite graph as relative papers demonstrated that noisy input data or representation improves the robustness of the neural network, and augmented the training dataset with different bipartite graphs with different distance cut-offs to relieve the protein-protein binding affinity data is not adequate, as we described in the A.1, less than 3k protein-protein binding affinity data available for ProAffinity++.

We augmented training data using different distance cut-offs when building the bipartite graph, in detail, the cut-offs mainly were 4 Å, 4.5Å, and 5Å for we found that the cut-offs between 4Å and 5Å resulted in a better performance of ProAffinity++(A.3.1).

The ablation result was shown in the table4. We found that adding noise significantly improved the correlation coefficient and led to an increase in MAE. However, the data augment only incurs slight movement, we attributed this slight improvement in performance to GNN is capable of obtaining bigger dilatation information from smaller dilatation through merging node features.

## 4 Methods

ProAffinity++ is a sequence- and structure-based protein-protein binding affinity predictor. Comparing these methods that predict the binding affinity based on the whole complex, we use the binding region structure as the evidence of predicting the binding affinity, and we use the many bipartite graphs to model the binding region of the protein complex, both sequence and structure level features were considered. We recognize that the binding affinity is influenced not only by the binding interface but also by the surrounding residues, which motivates our definition of the binding region as the residues from the binding interface and its surrounding residues. In this view, ProAffinity++ extracts high-level features from the binding region and encodes them into the hidden representation, then, the representation of the graph catted by physical energy term would be input into a multilayer perceptron(MLP) to predict the binding affinity.

We exhibit how to build a bipartite graph for a binary complex based on its binding region. While we build the bipartite graph from the complex structure, the input complex structure should be either an experimentally determined structure or a predicted structure, for now, protein structure predictors provide confident protein structure.

Once the complex structure is given, we determine the binding region according to the input complex structure. First, we calculate all distances of the residue pairs including inter-chain and intra-chain. Next, we identify residues from different chains in the binding interface with distances below the cutoff L1, and residues in the same chain that are directly in contact with the binding interface residues, for each chain separately. After the binding regions are determined, we build a bipartite graph that includes the node and the edge, every node of the bipartite graph is a residue in the binding region, and every edge means the relationship between the residue pair, the following residue pair will keep an edge, the residue pair in the binding interface with a distance lower than the cutoff L1 and from different chains, and the residue pair in the same chain with a distance lower than the cutoff L2.

To present the bipartite graph with abundant information we initialize the bipartite graph with interpretable node features and edge features. The node features include the residue evolutionary information embedded by the language model ESM-1v, the physicochemical properties of amino acids that are indexed by the AAindex, the structure information captured by the ESM-IF1, the physical energy term computed by FoldX with the AnalyseComplex module, the area of the interface computed using Pymol. The features details are shown in A.2.2. The edge features are composed of distance and the interaction type of the residues in the binding region. ProAffinity would update the representation of the bipartite graphs by a graph neural network, we used a message-passing neural network(MPNN) to update the information.

## 5 Discussion

ProAffinity++ is a novel protein-protein binding affinity predictor that addresses key limitations of existing methods. While traditional approaches are limited to predicting binding affinity for binary complexes and exhibit poor generalization, ProAffinity++ is capable of predicting binding affinity for multi-complexes by characterizing the entire complex and modeling all binding regions using bipartite graphs. This strategy reduces computational complexity by avoiding the need to model the structure of the entire complex.

ProAffinity++ outperforms other state-of-the-art protein-protein binding affinity prediction methods, and we attribute this superiority to two key factors. Firstly, a comprehensive set of protein complex data enables sufficiently training the ProAffinity++. Secondly, ProAffinity++ integrates diverse information from multiple sources to describe the protein complex, including sequence and structural features inferred by the pre-training model.

While ProAffinity++ demonstrates superior performance, there are still opportunities for further improvement. The first one is increasing the depth of the neural network layers, ProAffinity++ currently employs shallow MPNN to update the binding region representation and MLP to predict binding affinity, while deeper neural network layers possess better representation abilities suggesting that increasing the layer depth could lead to improved performance. Another one is augmenting the dataset with additional conformation, although ProAffinity++ leverages the binding state conformation of the complex, we believe that the conformation before binding also provides valuable information. By incorporating this additional conformational data into the dataset, we can potentially enhance the performance of ProAffinity++.

## Acknowledgments

We acknowledge the support from the National Key Research and Development Program of China (2024YFC3405501) and the National Natural Science Foundation of China (32271297). The Computer-X center at the Institute of Computing Technology supported the numerical calculations in this study. We also thank Dr. Haicang Zhang for helpful discussions.

## Declarations

## A Supplementary

### A.1 Dataset

The training set of ProAffinity++ including binary protein complex and multiprotein complex, and the performance of ProAffinity++ was tested on binary protein complex.

#### A.1.1 Binary protein complex

The training and validation set is the same as the split way of PPI-Affinity, including training set S653 and validation set S90.

#### A.1.2 Multiprotein complex

Multiprotein complexes come from PDBBind, we dropped the complexes with illegal PDB formats and the complexes that had many different binding affinities because different researchers measured those binding affinities with different devices and methods. Finally, we collected 1902 multiprotein complexes from the 2852 protein complexes, and we named this multiprotein complex dataset as S1902.

#### A.1.3 Nonsynonymous substitution dataset

The nonsynonymous substitution dataset includes the AT-Bind dataset and the Structural database of Kinetics and Energetics of Mutant Protein Interactions(SKEMPI).

The AT-Bind dataset is composed of 32 antibody-antigen complexes structures with 1101 mutations, and we select 645 complexes that only with single-point mutations as AT-Bind S1101 dataset.

The mutation data of the SKEMPI dataset is collected from the released paper, in this dataset, the protein binding affinity label may be lost or duplicated, we droped this data and finally got a dataset composed of 4005 mutations as SKEMPI S6706.

### A.2 Method

#### A.2.1 Loss Function

The loss function used by ProAffinity++ is HuberLoss. HuberLoss is a smooth loss function, where y is the true value, *ŷ*is the predicted value, and is a hyperparameter that controls whether the current loss function is square loss or absolute loss. When |*y* − *ŷ*|*< δ*, HuberLoss uses square loss, which can make the penalty for small errors relatively small, similar to the mean square error. When | *y* − *ŷ* |*> δ*, HuberLoss switches to absolute loss, which can make the penalty for large errors relatively small, thereby improving the robustness to outliers. This hyperparameter is selected through cross-validation. This feature of HuberLoss makes it more advantageous than the mean square error when dealing with some data with outliers or noise.

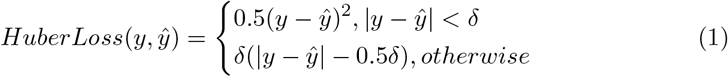

#### A.2.2 Features

**Table 5:** Summary of the node features of ProAffinity++.

| Feature | Tool | Dimension |
| --- | --- | --- |
| Physicochemical property of amino acid | AAindex | 57 |
| Evolutionary information | ESM-1v | 32 |
| Structure information | ESM-IF | 32 |
| Interface area | Pymol | 1 |
| Solution accessibility and secondary structure | DSSP | 7 |
| Sidechain rotation and oritation | PyRosetta | 9 |
| Physical energy term | FoldX | 21 |

The summary of the node features of ProAffinity++ is shown in the table5. We describe the features and the process method as follows:

1. The physicochemical property of amino acid is directly indexed by the AAindex to represent the amino acid chain identification, type, and physicochemical property.
2. The Evolutionary information captured by ESM-1v from the protein sequence and encoded into 1280 dimensions, to represent this evolutionary information more efficiently, the evolutionary information with 1280 dimensions is compressed into 32 dimensions by Principal Component Analysis(PCA).
3. Similar to the evolutionary information, the backbone structure information with 512 dimensions captured by ESM-IF is compressed into 32 dimensions by PCA. module of the FoldX.
4. The solution accessibility and secondary structure of residues are available through DSSP, furthermore, DSSP will output not only the secondary structure type but the properties of the secondary structure.
5. Sichain information included the sidechain barycentric coordinates relative to the C_*α*_ coordinates and the 6 rotation.
6. Physical energy term is the simulation and analysis result of FoldX based on force field to depict interaction of complex.

### A.3 Result

#### A.3.1 Graph type investigation

